# Assessment and refinement of sample preparation methods for deep and quantitative plant proteome profiling

**DOI:** 10.1101/273656

**Authors:** Gaoyuan Song, Polly Y Yingshan Hsu, Justin W. Walley

**Affiliations:** Department of Plant Pathology and Microbiology, Iowa State University, Ames, IA USA; Department of Biochemistry and Molecular Biology, Michigan State University, East Lansing, MI USA

## Abstract

A major challenge in the field of proteomics is obtaining high quality peptides for comprehensive proteome profiling by liquid chromatography mass spectrometry for many organisms. Here we evaluate and modify a range of sample preparation methods using photosynthetically active Arabidopsis leaf tissues from several developmental timepoints. We find that inclusion of FASP-based on filter digestion improves all protein extraction methods tested. Ultimately, we show that a detergent-free urea-FASP approach enables deep and robust quantification of leaf proteomes. For example, from 4-day-old leaf tissue we profiled up to 11,690 proteins from a single sample replicate. This method should be broadly applicable to researchers working on difficult to process samples from a range of plant and non-plant organisms.

**Abbreviations:**

Chloro: Methanol/Chloroform Extraction
FASP: Filter Aided Sample Prep
GO: Gene Ontology
IAA: Iodoacetamide
LFQ: Label Free Quantification
MS/MS: Tandem mass spectrometry
TF: Transcription Factor
UA: Urea Extraction
1D: 1 Dimensional
2D: 2 Dimensional

## Introduction

Proteins make or regulate nearly every component of a cell. Therefore, knowledge of the proteomic state of a cell and how it dynamically changes over time is critical to deepen our understanding of biological processes (1, 2). Further, deep coverage of the proteome enables proteogenomics projects that incorporate tandem mass spectrometry (MS/MS) proteomic data for genome annotation and refinement of gene models (3, 4). To achieve the goal of comprehensive and quantitative proteome profiling efficient protein isolation and digestion methodology is required.

Plant tissues present a significant challenge for isolating proteins and generating clean peptides suitable for peptide mass spectrometry. Plants produce relatively large amounts of interfering compounds such as phenolics, terpenes, pigments, organic acids, lipids, and polysaccharides. Furthermore, while interfering compounds are challenging in all plant tissues they are particularly abundant in green (i.e. photosynthetically active) tissues (5, 6). As a result numerous protein extraction methods using various combinations of trichloroacetic acid, acetone, methanol, chloroform, phenol, detergents, and molecular weight cutoff filters have been evaluated on plant tissues (5–22). However, nearly all the evaluations of extraction methods have examined compatibility with 2-dimensional gel electrophoresis rather than reverse phase liquid chromatography (LC), which is necessary for deep and quantitative proteome profiling.

We recently completed a large-scale protein atlas comprised of 33 different tissues generated from both vegetative and reproductive stages of development (23). Notably, profiling maize leaf tissue samples resulted in ∼31% fewer quantified proteins compared to non-photosynthetically active tissues. To address this discrepancy we examined if an improved sample processing protocol could enhance the depth of proteome coverage for photosynthetically active tissues. Here, we evaluate and optimize a range of protein extraction and digestion methods for proteome profiling using LC-MS/MS. In particular we focus on 1) urea extraction (UA), 2) methanol/chloroform extraction (Chloro), and 3) phenol extraction followed by in solution digestion. Additionally, we test each of these extraction methods coupled with a FASP-based (24) on filter digestion. Based on these initial analyses we used selected and tested a UA-FASP method for comprehensive 2D-LC-MS/MS profiling. Using this approach we are able to quantify up to 11,690 proteins from a single green-leaf sample.

## Experimental Procedures

### Plant Material

*Arabidopsis thaliana* accession Columbia (Col-0) was used for all experiments. Plants were grown on soil under 16 h/8 h (light/dark) and 22/20 °C (day/night) temperature. 21 days after planting rosette leaves were harvested and flash frozen in liquid nitrogen. For 4-day-old shoot samples, imbibed seeds were sown on a nylon mesh (Lab Pak; 250 µm) supported by a 1-inch high rack in Magenta vessel GA-7-3 (Sigma V8380) and grown hydroponically in liquid media (0.5X Murashige and Skoog salt, 1% sucrose, 0.5 g/L 2-(N-morpholino) ethanesulfonic, pH 5.7) shaken at 85 rpm under 16-hour light (∼110 µmol m^-2^ s^-1^) and 8-hour dark at 22°C. Shoot (herein referred to as leaf) tissue was harvested and flash frozen in liquid nitrogen. Prior to protein extraction leaf tissue was ground for 15 minutes under liquid nitrogen using a mortar and pestle.

### Protein Extraction and Digestion

#### Dilute SDS Extraction and Digestion

Digestion in dilute SDS following methanol/acetone extraction was performed as previously described (23, 25–28). Proteins were precipitated and washed with 50 ml of -20 °C methanol containing 0.2 mM Na_3_VO_4_ three times, then 50 ml of -20 °C acetone three times. Protein pellets were aliquoted into 2 ml Eppendorf tubes and dried in a vacuum centrifuge.

Protein pellets were suspended in 1 ml of 0.1% SDS, 1 mM EDTA, 50 mM Tris pH 7.5 at a 2:1 ratio of buffer:pellet and incubated at 94°C for 5 min. Protein concentration was measured using a Bradford assay (Thermo Scientific). Trypsin was added at a 1:100 ratio (enzyme:substrate) and the samples were digested at 37°C overnight. Iodoacetamide (IAA) was added to a final concentration of 12.5 mM and the samples were incubated at 37°C in dark for 15 min. The samples were centrifuged at 21,000 x g, for 5 min, the supernatant was transferred to a new tube, and 50 mM Tris pH 7.5 was added to the pellet. Following vortexing the samples were centrifuged at 21,000 × g, for 5 min and the supernatants were merged. Undigested protein was measured using Bradford assays. Then trypsin was added at a ratio 1:100 and LysC was added at 1/10 amount of trypsin. Digestion was performed at 37°C for 4 hrs. The samples were acidified to a pH of 2-3 using 100% formic acid and clarified by centrifugation at 21,000 × g for 20 min.The digested peptides were purified on a 150 mg Waters Oasis MCX cartridge to remove SDS. Peptides were eluted from the MCX column with 4 ml 50% isopropyl alcohol and 400 mM NH_4_HCO_3_ (pH 9.5) and then dried in a vacuum concentrator. Peptide amount was quantified using the Pierce BCA Protein assay kit with bovine serum albumn used to construct the standard curve.

#### SDS Extraction

The SDS-FASP protocol was slightly modified from Wiśniewski et al. (24) and performed as follows. 50 ml pre-chilled methanol was added to 0.25 g ground tissue, vortexed and kept at -20°C overnight. The samples were centrifuged at 4,400 x g for 15 min at 4 °C min. Next, 600 μl SDS buffer (4% SDS, 100 mM DTT) was added to the pellet. Samples were incubated at 95°C for 5 min while shaking in heated shaking block (Thermo). Next, the samples were sonicated 1 min with a probe sonicator at 70% intensity (Qsonica, Q125). This was repeated once with a 5 min heat incubation at 95°C with shaking. Samples then were clarified by centrifugation at 21,000 × g for 10 min, supernatant moved to a new tube, and centrifuged again a at 21,000 × g for 10 min. Protein yield was then measured using the Pierce 660 protein assay (Thermo Scientific). 2 mg of protein were then processed according to the FASP section below.

#### Urea Extraction

Lysis buffer (8M urea, 100mM Tris pH 7 and 5mM TCEP) was added at a ratio of 1:2 sample:buffer (w:v) to 0.25 g tissue. 1mm zirconium oxide beads (Next Advance) were added to the sample at ratio of 1:1 (v:v) and then the samples were shaken using a GenoGrinder at 1,500 rpm for 3 minutes. Samples were centrifuged at 4,000 × g for 3 min. The shaking and centrifuge steps were repeated once. Samples were transferred to a new tube and 4 volumes of prechilled 100% acetone was added. Samples were precipitated at -20°C for > 30 min followed by centrifugation at 4,500 × g for 10 min at 4°C. 80% acetone was added to the pellet and the sample was probe sonicated to resuspend the pellet and shear DNA. Samples were incubated - 20°C for > 5 min and then centrifuged at 4,500 × g for 10 min at 4°C. Precipitation and sonication in 80% acetone was repeated 3 times in total. Then prechilled 100% methanol was added to the pellet, sample was probe sonicated, and keep at -20°C for 30 min prior to centrifugation at 4,500 × g for 10 min at 4°C. Methanol precipitation was repeated once. During the final precipitation the sample was split in two, half of which was further processed by in solution digestion and half by FASP, prior to centrifugation. Finally, the samples were vacuum centrifuged till dry.

#### Methanol Chloroform Extraction

The methanol/Chloroform extraction protocol was based on a protocol described by Marx et al. (29). Extraction buffer (290mM sucrose, 250mM Tris pH8, 25mM EDTA pH8, and 10mM KCl) was added to 250 mg of ground tissue at a ratio of 5:1 (buffer:sample, v:w), vortexed, and then probe sonicated for 3 min. The samples were then filtered with miracloth. One volume of chloroform was added to the filtered extraction solution, the samples were vortexed, and then 3 volumes of H_2_O was added. Following vortexing the samples were centrifuged at 4,696 × g, for 5 min at 4°C. The resulting top layer was discarded, 3 volumes of prechilled methanol was added, the samples were vortexted to mix well, and then keep at -20°C for 2 hrs. The samples were then centrifuged at 4,696 × g for 5 min at 4°C, the supernatant was discarded, 4 volumes of prechilled 80% acetone was added, and then the sample was probe sonicated to resuspend the pellet. Samples were kept at -20°C overnight and then centrifuge at 4,500 × g, for 10min at 4°C. Precipitation with 80% acetone was repeated two more times with incubation at -20°C for 10 min. During the final precipitation the sample was split in two, half of which was further processed by in solution digestion and half by FASP, prior to centrifugation. Finally, the samples were vacuum centrifuged till dry.

#### Phenol Extraction

The phenol extraction protocol was modified from a protocol described by Slade et al., (30). 5 volumes (v:w) of Tris buffered phenol pH 8 was added to 250 mg of ground tissue, vortexed 1 min, then mixed with 5 volumes (buffer:tissue, v:w) of extraction buffer (50 mM Tris pH 7.5, 1mM EDTA pH 8, 0.9 M sucrose), and then centrifuge at 13,000 × g, for 10 min at 4°C. The phenol phase was transferred to a new tube and a second phenol extraction was performed on the aqueous phase. The two phenol phase extractions were combined and 5 volume of prechilled methanol with 0.1 M ammonium acetate was added. This was mixed well and keep at -80°C for 1h prior to centrifugation at 4,500 × g, for 10 min at 4°C. Precipitation with 0.1 M ammonium acetate in methanol was performed twice with incubation at -20°C for 30 min. The sample was resuspended in 70% methanol and split in two, half of which was further processed by in solution digestion and half by FASP, and precipitated. The supernatant was discarded and the samples were dried using a vacuum centrifuge.

#### In Solution Digestion

In solution digestion of proteins extracted using the UA, Chloro, or Phenol methods was performed as follows. Two volumes (buffer:pellet, v:v) of protein digestion buffer (8M urea, 50 mM Tris pH 7, 5 mM TCEP) was added to the pellet. The samples were then probe sonicated to aid in resuspension of the pellet. The protein concentration was then determined using the Bradford assay (Thermo Scientific). LysC was added with the enzyme to substrate ratio of 1:100 and then incubated at 37°C for 1h. Samples were then diluted 8-fold using 50 mM Tris pH 7 and trypsin was added with the enzyme to substrate ratio 1:50 prior to incubation at 37°C overnight. After the overnight trypsin digestion a Bradford assay was performed and a second trypsin digestion was carried out using an enzyme to substrate ratio of 1:100 for 4 hours at 37°C. IAA was added to the digested peptides at a final concentration of 12.5 mM and the samples were incubated at 37°C for 15 min in the dark. The samples were clarified twice by centrifugation at 21,000 × g for 20 min at room temperature. Finally, samples were desalted using 50 mg Sep-Pak C18 cartridges (Waters). Eluted peptides were dried using a vacuum centrifuge (Thermo) and resuspended in 0.1% formic acid. Peptide amount was quantified using the Pierce BCA Protein assay kit.

#### FASP-based on Filter Digestion

Protein was solubilized in 4 ml urea solution (8 M Urea, 0.1 M Tris-HCl pH 8), added to an Amicon Ultracel – 30K centrifugal filter (Cat # UFC803008), and centrifuged at 4,000 × g for 20-40 min. This step was repeated once. Then 4 ml of urea solution with 2mM TCEP was added to the filter unit and centrifuged at 4,000 × g for 20-40 min. Next, 2 ml IAA solution (50 mM IAA in 8 M urea) was added and and incubated without mixing at room temperature for 30 min in the dark prior to centrifuging at 4,000 × g for 20-40 min. 2 ml of urea solution was added to the filter unit, which was then centrifuged at 4,000 × g for 20-40 min. This step was repeated once. 2 ml of 0.05 M NH_4_HCO_3_ was added to the filter unit and centrifuged at 4,000 × g for 20-40 min. This step was repeated once. Then 2 ml 0.05M NH_4_HCO_3_ with trypsin (enzyme to protein ratio 1:100) was added. Samples were incubated at 37°C overnight. Then 1/5 original amount of trypsin (1ug/µl) and equal volume of Lys-C (0.1 µg/µl) were added and incubated for an additional 4 hours at 37°C. The filter unit was added to a new collection tube and centrifuged at 4,000 × g for 20-40 min. 1 ml 0.05M NH_4_HCO_3_ was added and centrifuged at 4,000 × g for 20-40 min. The samples were acidified to pH 2-3 with 100% formic acid and centrifuged at 21,000 × g for 20 min. Finally, samples were desalted using 50 mg Sep-Pak C18 cartridges (Waters). Eluted peptides were dried using a vacuum centrifuge (Thermo) and resuspended in 0.1% formic acid. Peptide amount was quantified using the Pierce BCA Protein assay kit.

### Liquid Chromatography

An Agilent 1260 quaternary HPLC was used to deliver a flow rate of ∼600 nL min^-1^ via a splitter for both 1D and 2D experiments. All columns were packed in house using a Next Advance pressure cell and the nanospray tips were fabricated using fused silica capillary that was pulled to a sharp tip using a laser puller (Sutter P-2000).

For 1D analyses 1 µg of peptides were loaded unto a 5 cm capillary column packed with 5 µM Zorbax SB-C18 (Agilent), which was connected to a 20 cm nanospray tip packed with 2.5 µM C18 (Waters) via a zero dead volume 1 µm filter (Upchurch, M548). Peptides were separated and delivered to the mass spectrometer using a 150 min reverse-phase gradient comprised of 5-30% (ACN, 0.1% formic acid) over 120 min, 30-80% (ACN, 0.1% formic acid) over 20 min, 80% (ACN, 0.1% formic acid) 5 min hold, and 80-0% (ACN, 0.1% formic acid) over 5 min.

For 2D analyses 30 µg of peptides were loaded unto a 20 cm capillary column packed with 5 µM Zorbax SB-C18 (Agilent), which was connected using a zero dead volume 1 µm filter (Upchurch, M548) to a 5 cm long strong cation exchange (SCX) column packed with 5 µm PolySulfoethyl (PolyLC). The SCX column was then connected to a 20 cm nanospray tip packed with 2.5 µM C18 (Waters). The 3 sections were joined and mounted on a custom electrospray source for on-line nested peptide elution. A new set of columns was used for every sample. Peptides were eluted from the loading column unto the SCX column using a 0 to 80% acetonitrile (ACN) gradient over 60 min. Peptides were then fractionated from the SCX column using a series of salt steps. For the initial test of the dilute SDS, SDS-FASP, and urea methods the following ammonium acetate salt steps were used: 10, 55, 60, 65, 70, 75, 80, 85, 90, 95, 100, 150, and 1000 mM. For the 2D analyses of 4-day-old and 3-week-old leaf tissue the following ammonium acetate salt steps were used: 10, 30, 32.5, 35, 37.5, 40, 42.5, 45, 50, 55, 65, 75, 85, 90, 95, 100, 150, and 1000 mM. For these analyses buffers A (99.9% H2O, 0.1% formic acid), B (99.9% ACN, 0.1% formic acid, C (100 mM ammonium acetate, 2% formic acid), and D (2 M ammonium acetate, 2% formic acid) were utilized. For each salt step a 150 min gradient program comprised of a 0-5 min increase to the specified ammonium acetate concentration, 5-10 min hold, 10-14 min at 100% buffer A, 15-120 min 5-35% buffer B, 120-140 min 35-80% buffer B, 140-145 min 80% buffer B, and 145-150 min buffer A was employed.

### Mass Spectrometry

The initial tests of the dilute SDS, SDS-FASP, and urea methods were carried out using a Thermo Scientific Q-Exactive high-resolution quadrupole Orbitrap mass spectrometer. Data dependent acquisition was obtained using Xcalibur 3.0.63 software in positive ion mode with a spray voltage of 2.00 kV, a capillary temperature of 275 °C, and a RF of 70. MS1 spectra were measured at a resolution of 70,000, an automatic gain control (AGC) of 3e6 with a maximum ion time of 100 ms and a mass range of 400-2000 m/z. Up to 15 MS2 were triggered at a resolution of 17,500. An AGC of 1e5 with a maximum ion time of 50 ms, an isolation window of 4.0 m/z, and a normalized collision energy of 28 were used. Charge exclusion was set to 5-8, >8. MS1 that triggered MS2 scans were dynamically excluded for 15 s.

The UA, UA-FASP, Chloro, Chloro-FASP, Phenol, and Phenol-FASP samples were analyzed using a Thermo Scientific Q-Exactive Plus high-resolution quadrupole Orbitrap mass spectrometer. Data dependent acquisition was obtained using Xcalibur 4.0 software in positive ion mode with a spray voltage of 2.00 kV and a capillary temperature of 275 °C and an RF of 60. MS1 spectra were measured at a resolution of 70,000, an automatic gain control (AGC) of 3e6 with a maximum ion time of 100 ms and a mass range of 400-2000 m/z. Up to 15 MS2 were triggered at a resolution of 17,500. An AGC of 1e5 with a maximum ion time of 50 ms, an isolation window of 1.5 m/z, and a normalized collision energy of 28 were used. Charge exclusion was set to unassigned, 1, 5-8, and > 8. MS1 that triggered MS2 scans were dynamically excluded for 15 s (2D analysis of 3-week-old UA-FASP samples) or 25 s (1D analyses and 2D 4-day-old samples).

### Data Analysis

The raw data were analyzed using MaxQuant version 1.6.0.16 (31). Spectra were searched against the Tair 10 proteome, which was complemented with common contaminants by MaxQuant. Carbamidomethyl cysteine was set as a fixed modification while methionine oxidation and protein N-terminal acetylation were set as variable modifications. Digestion parameters were set to “specific” and “Trypsin/P;LysC”. Up to two missed cleavages were allowed. A false discovery rate less than 0.01 at both the peptide spectral match and protein identification level was required. The ‘second peptide’ option was on to identify co-fragmented peptides. The match between runs feature of MaxQuant was not utilized. Quantification was performed using the MaxQuant label-free quantification (LFQ) algorithm (32) or MS/MS count (spectral counting) output. Scatterplots and Pearson correlation values were generated using Perseus (33). Gene Ontology enrichment analysis was performed using GOrilla http://cbl-gorilla.cs.technion.ac.il/ (34). The list of Arabidopsis transcription factors was obtained from a curated set of 2,492 transcription factors reported by Pruneda-Paz et al. (35).

## Results and Discussion

We initially tested three protein extraction/digestion methods using 2D-LC-MS/MS. Specifically, we examined 1) a dilute SDS in solution digestion method we have used extensively (23, 25–28), 2) SDS-FASP (24), and 3) a urea extraction and in solution digestion method (UA). We found that the UA method yielded the highest number of detected proteins (Supplemental Table 1). This observation maybe due to incomplete removal of SDS from the other two methods, which has been previously reported for SDS-FASP (36). Thus, we next tested several SDS-free sample processing approaches.

**Table 1.** Comparison of protein recovered (µg) using chloroform, phenol, and urea based extraction methods from 100 mg of 3-week-old Arabidopsis leaf tissue. Protein amount was quantified using Bradford assays. Data are means of n=3 ± SEM.

| Chloroform | Phenol | Urea |
| --- | --- | --- |
| $279.8 \pm 17.6$ | $908.2 \pm 19.5$ | $1004.7 \pm 23.7$ |

Based on the initial performance of the UA method we decided to test it in further detail and also compare it to methanol/chloroform (Chloro) and phenol based methods. First, we examined the amount of protein that was recovered from these extraction approaches. For this we performed extractions from 100 mg of green leaf tissue from 3-week-old Arabidopsis plants and then quantified the amount of protein recovered, prior to in solution digestion, using Bradford assays. The UA method yielded, on average, 1,004.7 µg of protein, which was the largest amount of protein recovered (Table 1). Additionally, both the phenol and UA performed significantly better than the Chloro method in terms of amount of protein recovered. Therefore, in situations where there is limited starting material the phenol and UA methods have a distinct advantage.

Next, we evaluated these methods using mass spectrometry. Furthermore, because molecular weight cutoff filters should offer additional removal of non-protein contaminates we examined whether the performance of UA, Chloro, or phenol methods was improved with inclusion of FASP-based on filter digestion (Supplemental Figure 1). We analyzed triplicate technical replicate extractions for each method using 1 µg of peptides by 1D-LC-MS/MS (Figure 1 and Supplemental Table 2). The resulting data exhibited high quantitative reproducibility based on Pearson correlation values of MaxQuant LFQ intensity among the replicates of each extraction method (Supplemental Figures 2-7). All extraction methods were improved by addition of FASP-based on filter digestion (Figure 1). Specifically, FASP resulted in increases in the number of protein identified of 10.2%, 7.1% and 12.8% for the Chloro-FASP, phenol-FASP, and UA-FASP, respectively. In particular, the increase due to FASP was evident at the MS/MS level where it resulted in a 20.7%, 15.3%, and 44.6% increase for Chloro-FASP, phenol-FASP, and UA-FASP, respectively. Finally, in terms of proteins detected, unique peptides detected, and MS/MS identified the phenol-FASP and UA-FASP methods perform similarly to one another and both outperform the Chloro-FASP method (Figure 1).

**Figure 1.**
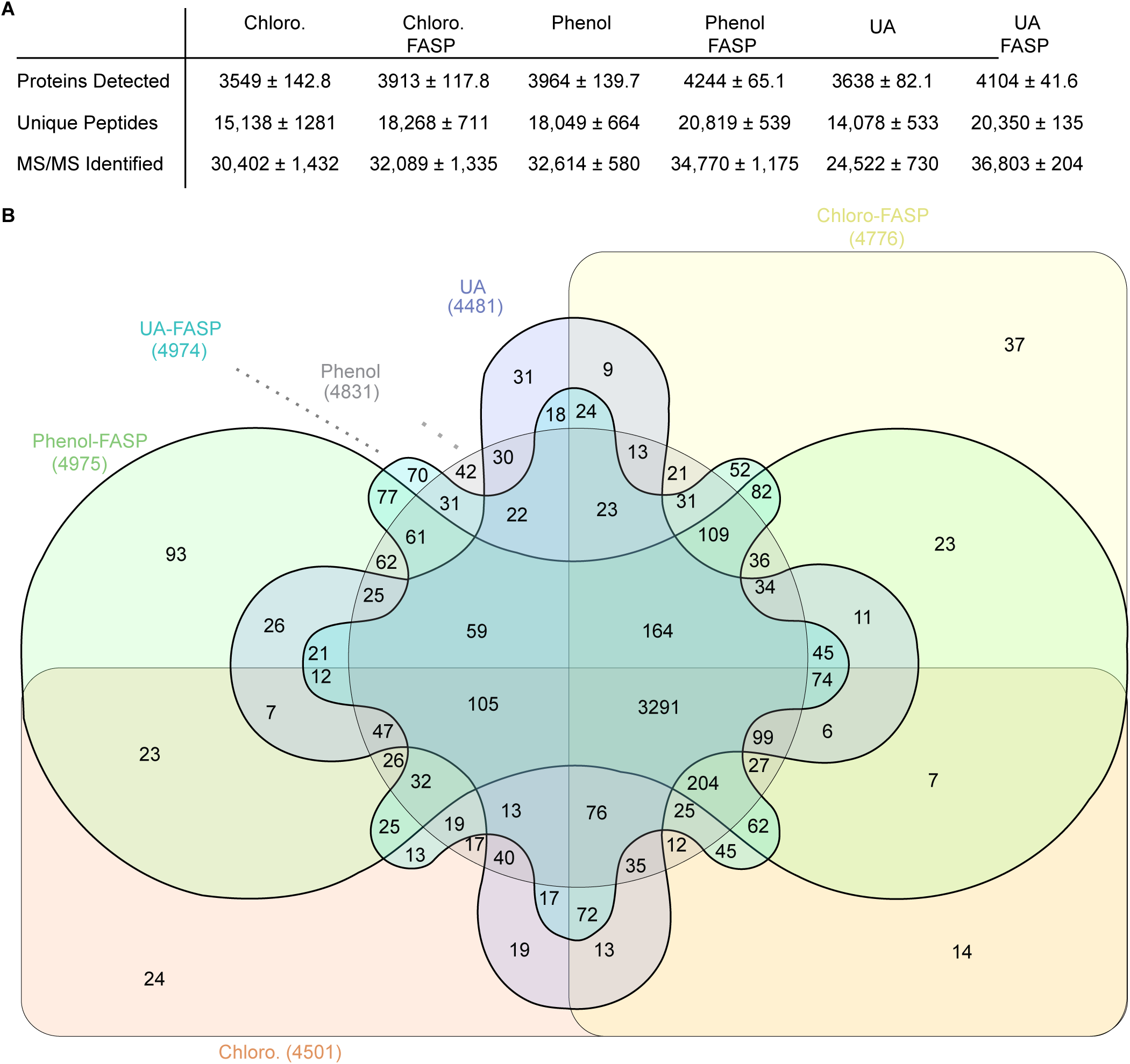
Evaluation of the performance of sample processing methods for LC-MS/MS. **A)** Technical replicate extractions were performed in triplicate on the same starting tissue for each method. Data are means of 3 ± SEM. **B)** Overlap in the number detected proteins between the sample processing methods. Data are the total identified proteins from triplicate runs.

**Figure 2.**
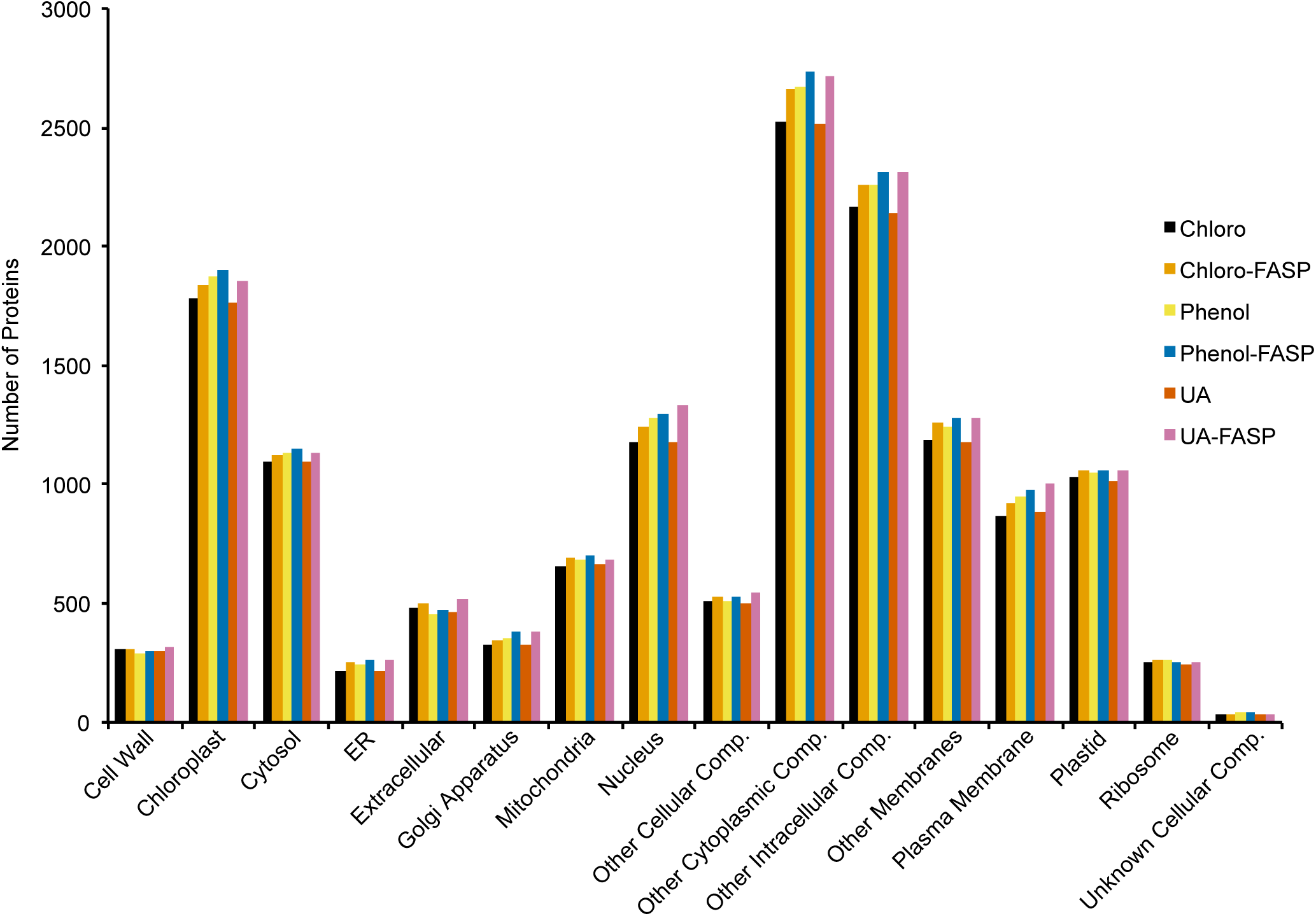
Comparison of GO cellular component. Total number of proteins per subcellular compartment identified for each sample processing method

We examined the subcellular localization of the proteins detected by each method using Gene Ontology (GO) cellular component annotations. This analysis revealed no major biases in the localization profile of the proteins extracted from each method (Figure 2). Furthermore, we performed GO term enrichment analysis on the proteins specifically detected by each method. This analysis did not return enrichment for any subcellular “component” GO term for any of the sample preparation methods. Based on these data it does not appear that any of the methods we tested exhibit a subcellular compartment extraction bias relative to the other tested methods.

Encouraged by these results we sought to test the performance of UA-FASP processed peptides for deep proteome profiling by employing online salt fractionation coupled with mass spectrometry. For this experiment we performed UA-FASP in duplicate by extracting protein from independent biological replicates of 3-week-old Arabidopsis leaf tissue. Significantly, this analysis revealed that the UA-FASP method enables deep profiling of an average of 10,242 proteins, 76,241 unique peptides, and 448,988 MS/MS per leaf tissue sample (Figure 3 A-C and Supplemental Table 3). Furthermore, we examined reproducibility of the biological replicate analyses and found high Pearson correlation values for quantification using either MaxQuant LFQ intensity (*r* = 0.966) or spectral counting (*r* = 0.938) (Figure 3 D&E).

**Figure 3.**
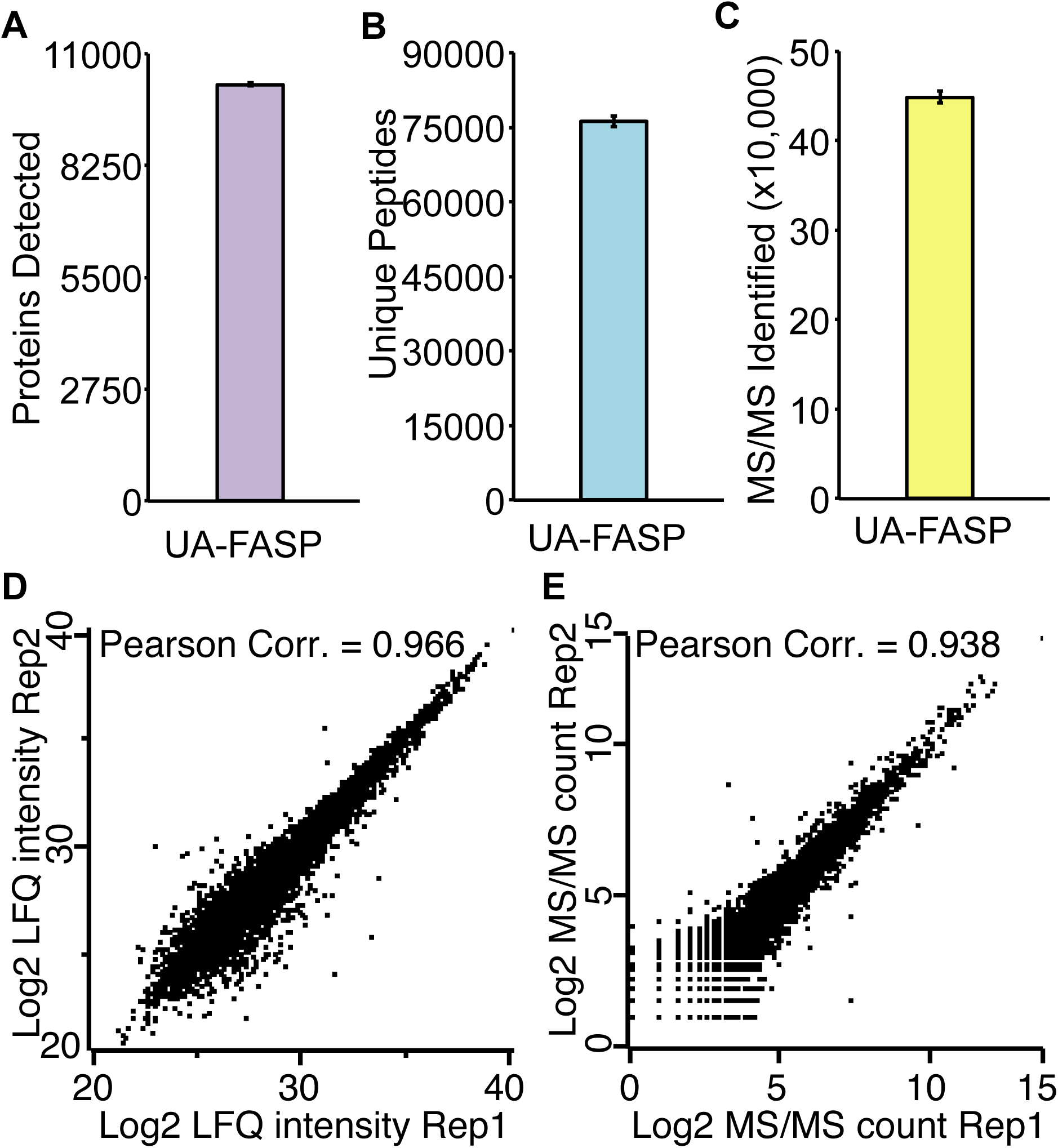
UA-FASP enables deep profiling of leaf tissue. Proteins were extracted from three-week-old Arabidopsis leaf tissue and processed into peptides using the UA-FASP method. **A-C)** Data are means of two independent biological replicates ± SEM. Scatterplots of the two biological replicates where the proteins are quantified as **D)** LFQ intensity or **E)** spectral counts.

The composition of leaves changes during development, which may impact sample preparation performance. Therefore, we next tested the UA-FASP method using 4-day-old leaf tissue. 2D-LC-MS/MS was performed similarly to the 3-week-old tissue with the only difference being that we extended the dynamic exclusion setting from 15 to 25 seconds. We identified up to 11,690 proteins from a single sample and an average of 11,055 proteins, 89,298 unique peptides, and 415,596 MS/MS from the two biological replicates (Supplemental Tables 4&5). The run-to-run reproducibility between biological replicates was also high with Pearson correlation values of 0.95 (LFQ) and 0.98 (spectral counting) (Supplemental Table 5). Taken together these data reveal that the UA-FASP method enables deep and reproducible proteome quantification.

Finally, to highlight the depth of proteome coverage enabled using UA-FASP coupled with 2D fractionation we examined the detection of transcription factors (TFs), which are typically low abundance proteins. We were able to detect 576 TFs in 3-week-old leaves and 669 TFs in the 4-day-old leaves (Supplemental Tables 3&4). Furthermore, the detected TFs arose from a broad range of TF families (Figure 4), which suggests that the high TF coverage we observe is not due to detecting a large number of TFs from just a few high abundance TF families. Taken together these data reveal that the UA-FASP method enables deep and reproducible proteome quantification.

**Figure 4.**
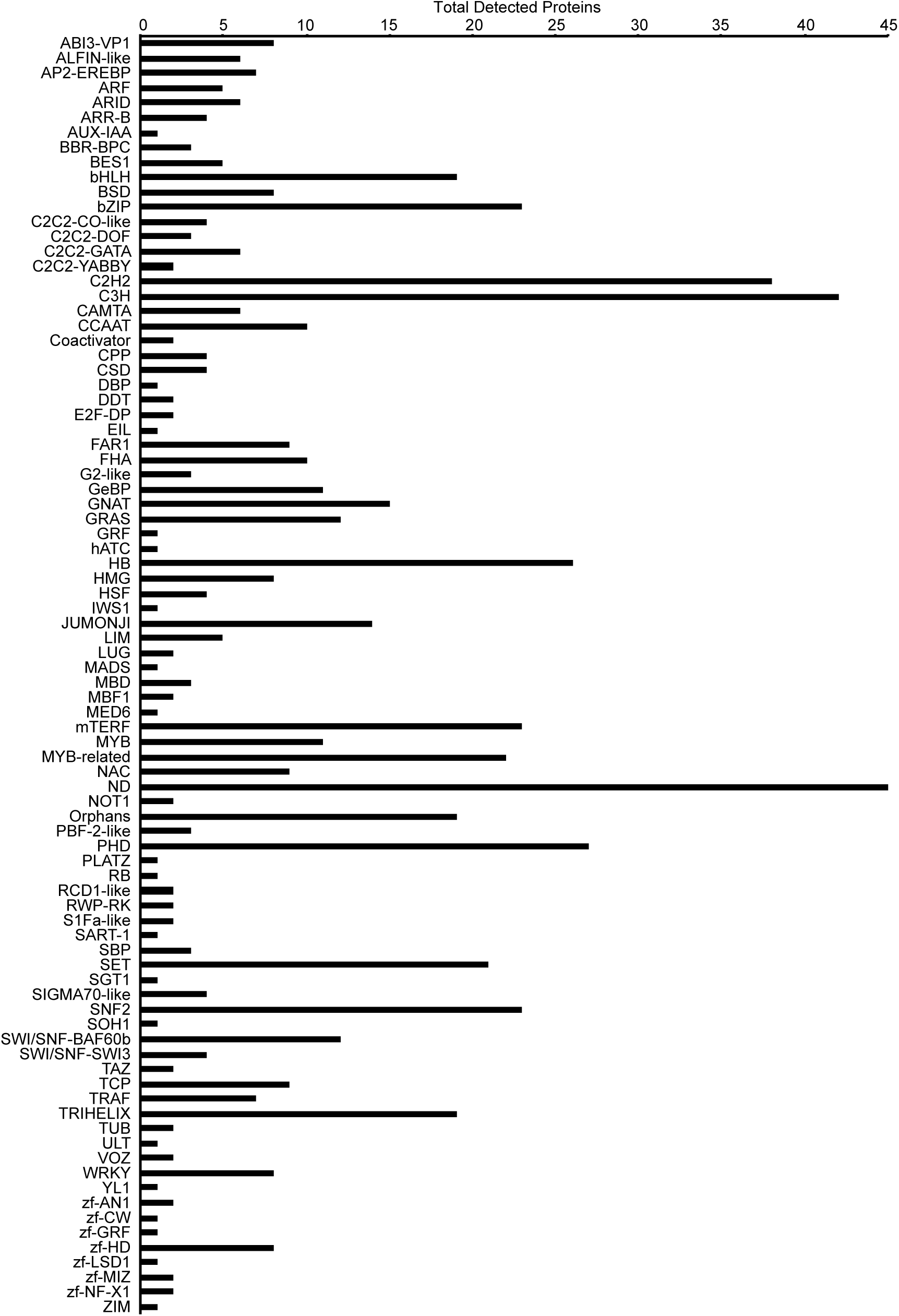
Summary of transcription factors identified. Total number of transcription factors identified per family using 2D-LC-MS/MS of UA-FASP processed four-day-old leaf tissue.

## Concluding Remarks

Obtaining clean peptides for comprehensive proteome profiling remains challenging. This is especially the case for difficult samples such as photosynthetically active green leaf tissues. While there are many sample extraction and processing methods that have been developed most evaluations of their efficacy on plant tissues has been carried out using 2-dimensional gel electrophoresis rather than reverse phase LC prior to mass spectrometry. To address this we evaluated several approaches for sample processing that are based on popular urea extraction, methanol/chloroform, and phenol methods. Additionally, we examined whether these methods were improved using FASP-based on filter digestion. We found that phenol-FASP and UA-FASP methods enabled the best coverage of protein, peptides, and MS/MS based on 1D-LC-MS/MS analyses. We chose the UA-FASP method for further testing because 1) it resulted in the largest recovery of protein (Table 1), 2) enabled detection of a similar total number of peptides and proteins as the phenol-FASP method (Figure 1), and 3) it does not require the use of hazardous phenol. Furthermore, because the UA-FASP method does not require complicated phase separation it should be compatible with recently described approaches for high-throughput sample preparation using 96-well filter plates (37, 38). Ultimately, using the UA-FASP method we were able to robustly and reproducibly quantify over ten thousand proteins from each analyzed leaf sample. We believe this method will enable comprehensive profiling of proteomes from difficult to process samples from a range of plant and non-plant organisms.

## Data Availability

The raw spectra for the proteome data have been deposited in the Mass Spectrometry Interactive Virtual Environment (MassIVE) repository: https://massive.ucsd.edu/ProteoSAFe/static/massive.jsp with the MassIVE dataset identifier MSV000082080.

## Acknowledgments

This work was supported by NIH grant 1R01GM120316-01A1, United States Department of Agriculture National Institute of Food and Agriculture Hatch project 3808, and by the Iowa State University Plant Science Institute (PSI).

